# Molecular dynamics simulations reveal the impact of NUTD15 variants in structural conformation and dynamics

**DOI:** 10.1101/2022.10.23.513377

**Authors:** Elena Gómez-Rubio, Javier Garcia-Marin

## Abstract

NUDT15 or MTH2 is a member of NUDIX protein family that catalyze the hydrolysis of nucleotides and deoxynucleotides, including thioguanine analogues. NUTD15 has been reported as a DNA sanitizer in humans, and more recent studies have proved that genomic variants are related to a poor prognosis in inmoplastic and immunologic diseases with thioguanines. Despite of this, the role of NUTD15 in physiology and molecular biology is quite unclear, as well as the mechanism of action of this enzyme. The existence of clinically relevant variants has prompted the study of this enzymes, whose capacity to bound and hydrolyze thioguanine nucleotides is still poor understood. By using a combination of biomolecular modelling techniques together with molecular dynamics we have studied the monomeric wild type NUTD15, as well as two important variants R139C and R139H. Our findings reveal not only how nucleotide binding stabilizes the enzyme, but also how two loops are responsible for keeping the enzyme in a packed close conformation. Mutations in α2 helix affect a network of hydrophobic and π-interactions that are responsible of active site enclosing.

## 1 Introduction

In living organisms, nucleic acids are macromolecules whose fundamental building blocks, known as nucleotides, are susceptible to a variety of chemical modifications. Some of these transformations are due to physiological circumstances such as post-translational or epigenetic modifications, which regulate and tune the cell cycle (Liyanage and others 2014). However, chemical modifications of DNA can also be due to non-enzymatic processes including oxidative stress, ionizing radiation or radiotherapy (Belli and Tabocchini 2020; Lomax, Folkes, and O’Neill 2013; Swenberg and others 2010). In this regard, the human body must have some kind of mechanism to repair DNA nucleobase alterations either, when they are forming part of DNA double strand or as unprotected nucleotides in solution, which are more susceptible to chemical alterations (Topal and Baker 1982).

The NUDT15, also known as the nudix hydrolase 15 or MTH2, is an enzyme that belongs to the NUDIX (nucleoside diphosphates linked to a moiety x) superfamily of proteins (Maki and Sekiguchi 1992). It was first identified in *Escherichia coli* as a member of the sanitizer pool of enzymes responsible for detoxification due to oxidized nucleotides accumulation. This protein catalyzes the hydrolysis of canonical nucleotides, as well as oxidized deoxynucleotide triphosphates (dNTPs), including substrates like 8-oxo-dGTP, whose incorporation into DNA can lead to mispairing among base pairs, and thus to insidious consequences.

NUDIXs substrate specificity has been the object of extensive research, and recent studies have proved that NUTD15 is able to hydrolyze metabolites of thiopurines drugs like 6-thio-deoxyguanosine triphosphate (Rehling and others 2021; Valerie and others 2016). Azathioprine, mercaptopurine and thioguanine belong to the group of thiopurine antimetabolites, a kind of drugs that, after their bioactivation inside the cell into thioguanosine triphosphate (TGTP), inhibit several enzymes belonging to the purine biosynthesis and produce misincorporation into DNA and RNA (Figure 1) (Karran and Attard 2008). Thiopurine drugs are used as anticancer and immunosuppressive agents in the treatment of different malignances, including lymphoblastic leukemia, acute myeloid leukemia, and inflammatory bowel disease (Cargnin and others 2018; Terdiman and others 2013; Pui, Robison, and Look 2008).

**Figure 1.**
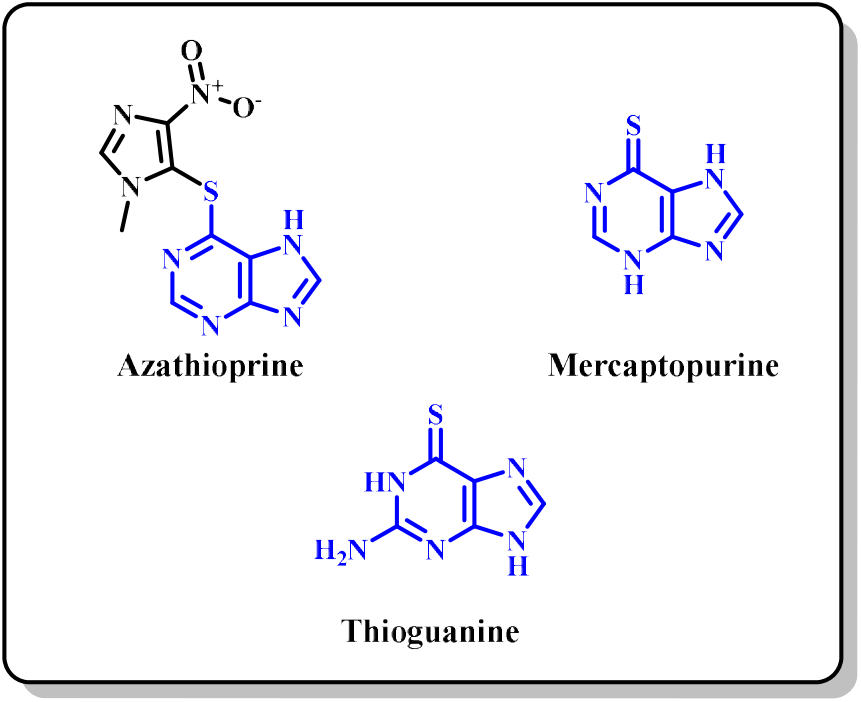
Molecular structures of thiopurine drugs approved for the treatment of different diseases. Highlighted in blue the pharmacophoric scaffold of thiopurine.

Several clinical investigations, including a genome-wide association study, have recently described a missense variant in the NUDT15 gene rs116855232 (Arg139 substitution by Cys), which leads to the accumulation of 6-mercaptopurine metabolites and cellular toxicity (originating thiopurine-induced leukopenia) during thiopurine treatments (Yang and others 2014; Singh and others 2017).

In this context, the interest of the scientific community in the research of NUTD15, its role in purine metabolism and biology has increased during the last few years. Recently, an active site targeting chemical probe was developed to study the protein as a potential drug for thiopurine combination therapy (Zhang and others 2020).

However, there are still a lot of questions that remain elusive for NUTD15 such as how it recognizes substrates, the catalytic mechanism, or the impact of point mutations on the activity of clinically relevant variants. To answer some of these questions, a detailed view of the dynamic behavior of the protein at the atomic level is needed. To address some of those unknowns concerning to NUTD15 clinical variants, we have performed exhaustive molecular dynamics simulations (MD) in order to shed some light on the conformational changes that the protein undergoes. We compare the wild type (WT) NUTD15 with some of their most important variants in the bound and unbound state. This knowledge will facilitate the understanding of NUTD15 genetic variants, as well as the design of more effective drugs or even the optimization of this enzyme as a biocatalyst.

## 2 Material and methods

### 2.1 Protein preparation

A model of NUDT15 with the whole sequence (residues 1-164) was built using the Robetta web server (https://robetta.bakerlab.org/). For this purpose, the RosettaCM protocol was selected, choosing a single template method and considering PDB: 6T5J as a starting coordinates, to generate a single model (Song and others 2013). The RosettaCM protocol allows building models by homology modeling in those regions for which there is sequence alignment, while those for which there is no alignment are built using a *de novo* method. In the case of NUDT15, it has applied homology modeling for residues 10 to 163, and it was generated *de novo* the fragment corresponding to residues 1 to 9 and residue 164. The quality of the model was characterized using the QMEAN web server, applying the QMEANDisCo scoring function (Studer and others 2020). The evaluation of the model resulted in a Qmean score of 0.84 ± 0.07, indicating good quality.

Protonation states of NUTD15 titratable residues were assigned using the ProteinPrepare module available at the https://www.playmolecule.com website (Martínez-Rosell, Giorgino, and De Fabritiis 2017). All histidines were considered in their delta tautomeric form except His25, His49 and His91 which were protonated at the epsilon position. To generate TGTP complexes, 6-thio-guanosine triphosphate was manually built up starting from PDB: 5LPG, which contains a molecule of 6-Thio-guanosine monophosphate in the active site (Valerie and others 2016). The molecular graphics program PyMOL 2.1 (The PyMOL Molecular Graphics System, Version 2.0 Schrödinger, LLC.) was employed for completing the missing β and γ phosphate groups of the nucleotide. NUTD15 variants were generated by means of CHARMM-GUI interface during the protein preparation step using the mutation tool (Jo and others 2008).

### 2.2 Molecular dynamics simulation

The systems were built using the Solution Builder tool from the CHARMM-GUI server (https://www.charmm-gui.org/). Each protein or protein-ligand complex was solvated into a cubic box extended 8 Å from solute, then Na^+^ and Cl^-^ ions were added to neutralize the systems. Water molecules were treated with the TIP3P model, AMBER ff14SB force field was applied for proteins (Maier and others 2015), and parameters for the TGTP molecule were generated using the CHARMM general force field (CGenFF2) (Wang and others 2004).

Energy minimization was performed via the steepest-descent method during 5000 steps, with a tolerance of 1000 kJ mol^-1^nm^-1^. The van der Waals forces were defined with a cutoff of 9 Å, while the cutoff for the particle mesh Ewald (PME) method applied to consider the long-range electrostatic interactions was set on 9 Å (Essmann and others 1995). Long range dispersion corrections were applied for energy and pressure. The neighbor search was carried out with the Verlet cut-off scheme, being updated every 10 steps. For all hydrogen-containing bonds, harmonic constraints were applied using the LINCS algorithm. For the backbone of the protein and the protein side chains, harmonic potential restraints of 400 kJ mol^-1^ nm^-1^ and 40 kJ mol^-1^ nm^-1^ were applied, respectively. The equilibration phase was performed at the canonical ensemble (NVT) and was carried out during 125 ps. In this stage, for temperature coupling, a Noose-Hover extended ensemble was used, with a bath temperature of 298 K and a time constant of 1 fs. The cut-offs for van der Waals forces and long-range electrostatic interactions were treated as in the minimization step, but the Verlet pair list was updated every 20 steps. The Maxwell distribution, starting from a pseudo random seed, was used to generate the initial velocities. Finally, the production phase was carried out using a time step of 2 fs, at the NTP ensemble controlling the temperature with the Nose-Hoover thermostat to maintain the system at 298 K and maintaining a constant pressure of 1 bar using the Parrinello-Rahman barostat. All simulations were conducted using Gromacs 2020.21 on a Nvidia GeForce GTX 980, resulting in an aggregate simulation time of 3 μs.(Bauer, Hess, and Lindahl 2022).

### 2.3 Analysis

For analysis purposes, all trajectories were centered on the protein atoms and aligned to the first frame using Gromacs. RMSD, RMSF and distances were measured using MDtraj (McGibbon and others 2015) and VMD (Humphrey, Dalke, and Schulten 1996) softwares. After production runs, the RMSD of each independent simulation was used to discard the data in the equilibration phase, so mean RMSD quantities were computed after discarding the initial 75 ns of production run. Protein-ligand contacts were calculated on the whole trajectory, setting a threshold of 3.5 Å. Occupancy volumetric maps were generated via the VolMap plugin implemented in VMD. Hydrogen bonds were defined by acceptor-donor atom distances shorter than 3.0 Å. Statistical analysis between data sets was carried out by means of t-test via SciPy library implementation in Python 3.7 (Virtanen and others 2020), considering a significance threshold p < 0.05. The visualization tool VMD was used to visualize trajectories and, together with ChimeraX (Pettersen and others 2021), to render all pictures, while graphics were generated with the Matplotlib 3.1.0 library (Hunter 2007).

## 3 Results and discussion

### 3.1 Protein stability and dynamics

Among the different NUTD15 variants identified up to date, we selected for our purposes two well characterized mutants, both with substitutions in the position of Arg139 by a cysteine (R139C) and histidine (R139H). This decision was based in the clinical relevance of these mutants in thiopurine therapy and the availability of experimental data to validate and compare our computational results (Table 1) (Yang and others 2014; Rehling and others 2021; Cargnin and others 2018).

**Table 1.** Enzymatic parameters of NUTD15 described in the literature (Rehling and others 2021).

| Enzyme 1<sup>a</sup> | WT | R139C | R139H |
| --- | --- | --- | --- |
| T<sub>m</sub>(°C) | 58 | 50 | 52 |
| T<sub>m</sub>(°C)<sup>a</sup> | 63 | 56 | 57 |
| Normalised activity | 100 | 30 | 18 |

<sup>a</sup> In presence of TGTP.

NUTD15 has been crystallized in form of homodimeric complexes with or without inhibitors or nucleotides (Carter and others 2015; Carreras-Puigvert and others 2017; Rehling and others 2021; Valerie and others 2016; Zhang and others 2020). Due to the total symmetry of the crystallographic assembly, we decided to perform MD simulations using the monomeric form of NUTD15. Since the *N*- and *C*-termini motifs of the enzyme present a high level of disorder and they have not been solved, homology modeling techniques were used to build a whole structure of NUTD15 spanning residues 1-164 (Uniprot code: Q9NV35).

The model was generated by RosettaCM presenting a high overlap with the crystallographic structure of NUDT15 used as a template (PDB 6T5J) (Song and others 2013), with an RMSD of 0.48 Å calculated on the alpha carbon trace. The main differences are located in the unstructured regions corresponding to residues Val101-Pro109 and Pro148-Leu156, with an RMSD of 0.785 and 1.995 Å respectively. Regarding the active site, minimum changes are observed, bending approximately 90° with respect to the initial crystal coordinates in the orientation of the side chains of Gln44, His49, Tyr90 and Tyr92.

Once a plausible model of NUTD15 was generated, to assess its feasibility, we simulated apo and holo (in complex with TGTP) NUTD15 structures over 500 ns of unrestricted classical MD (supporting info. Figure S1). Simulations of the explicitly solvated system revealed different results when comparing apo to holo proteins. Trajectories converged after approximately 100 ns of production dynamics, either for apo and holo complexes (supporting info. Figure S1). These findings proves that the monomeric form of NUTD15 is stable when simulated in the presence of explicit water and/or in the presence of this nucleotide, and also suggest that this could be extended to other cognate ligands. When no TGTP was bound in R139C mutant, some noticeable fluctuations were observed in the protein *N*-tail, which corresponded to an outward-inward movement of this motif from the initial position (supporting info. Figure S2). This segment of the protein would be in close contact with a second monomer of the NUTD15 homodimeric assembly (PDB: 6T5J) (Zhang and others 2020), used as a template for our homology models. However, after these fluctuations, the *N*-termini went back to the initial position. Since crystal structures did not solve this protein section due to high mobility and disorder structure, our observations are in line with experimental results (Carter and others 2015; Carreras-Puigvert and others 2017; Rehling and others 2021). Additionally, this observation proves that the monomeric enzyme can exist in a similar three-dimensional configuration compared to the crystalized homodimeric assemblies, which is also observed *in vivo* (Rehling and others 2021).

To further study protein dynamics and obtain deeper insights into the behavior of NUTD15, we also calculated the mean RMSD value across the simulation (Figure 2). Some qualitative correlations could be found among apo values and protein folding temperature (Tm, see Table 1). While apo WT displayed a mean RMSD of 2.27 ± 0.13 Å, substitution of Arg139 by a cysteine leads to a significant increment of 2.49±0.26 Å (p<0.005), according to a less stabilized system during MD simulations. This is in good agreement with experimental stability data measured in terms of Tm (see Table 1), which accounts for the protein stability with a decrease in Tm for this variant. Interestingly, this tendency is not observed in the R149H mutant, whose Tm is lower compared with the wild type (WT). Regarding protein-ligand complexes, it was found that the presence of TGTP molecule in the catalytic site produces a slight stabilization of NUTD15, as illustrated by mean RMSD values lower when compared with apo states in all the cases (Figure 1A).

**Figure 2.**
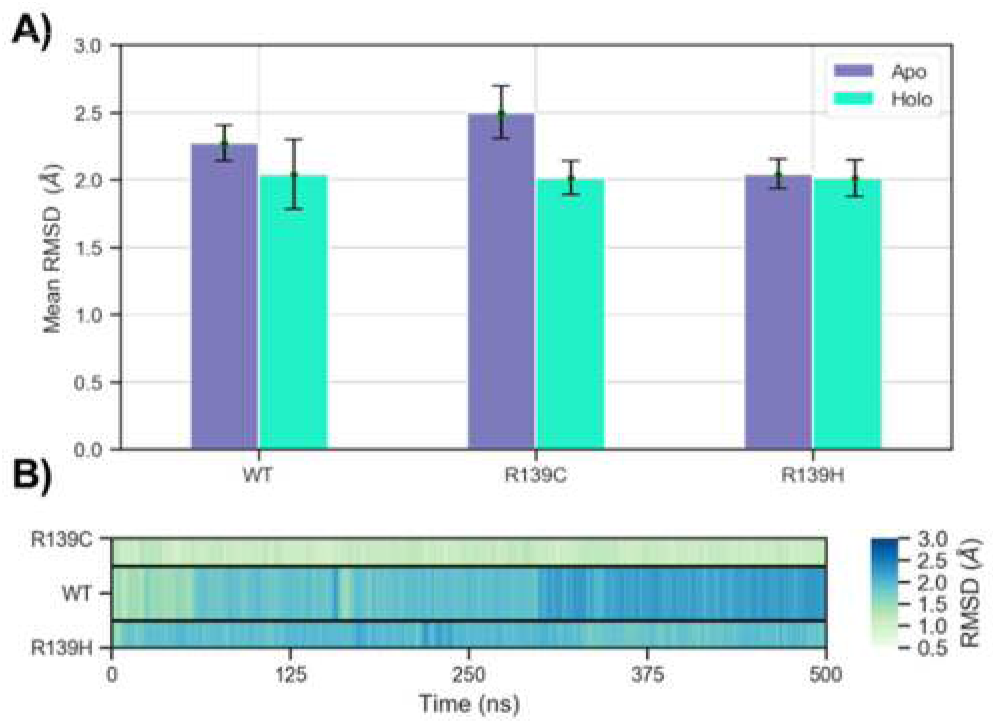
A) Barplot of mean RMSD values ± Standard Deviation (SD) for backbone atoms in all apo and holo MD simulations. B) Heatmap plot of the RMSD values for TGTP heavy atoms over time.

This effect has been previously observed in other computational studies which consider this metric to study the effect of direct ligand binding when a molecule acts towards the active site of the enzyme (García-Marín, Rodríguez-Puyol, and Vaquero 2022; Makurat, Cournia, and Rak 2022; Ge and others 2017). This tendency is also in accordance with Tm values when the inhibitor is present in the assay, confirming the ability of the substrate to stabilize protein structure (Rehling and others 2021; Man and others 2019). Although this situation is also observed in the R139H variant, the subtle change seems to be not quite significant (p=0.66) for this variant, which is the least catalytically active. All in all, this data confirms the suitability of our computational biomolecular models to study NUTD15 systems and also confirms that TGTP can stabilize not only the surroundings of the active site, as previously reported, but the whole protein instead (Man and others 2019).

The lack of structural information about protein-substrate dynamics and interactions prompted us to carry out MD studies, including the nucleotide 6-thio-guanosine triphosphate (TGTP) bound in WT, R139C, and R139H NUTD15. First, a thorough analysis of the cognate nucleotide inside the active site pocket was carried out in order to check possible differences in ligand dynamic binding mode. In these holo complexes, only minor conformational changes during simulations were observed, indicating that the ligand binding pose is extremely stable, with RMSD values below 3 Å (Figure 1B). In general, only one metaestable binding mode was observed during our simulations, coincident we that observed in the crystallographic structure.

The TGTP molecule does not suffer great shifts or movements during our simulations in either WT or mutant variants. In fact, ligand remains firmly anchored to the active site thanks to two hydrogen bonds between the purine amino group with Leu45 and Val16 carbonyls. This binding mode is reinforced by an extra hydrogen bond between the pyrimidine N1 nitrogen and carboxamide -NH2 from Gln44. Sulfur atom is deep buried into a small subpacket where interacts with -NH from Leu45 and -Leu138. A strong electrostatic interaction is provided by Arg34 which binds the α phosphate from TGTP thanks to its negatively charged oxygen atom. The main interactions are also complemented with CH-π interactions from Phe135 and Phe136 towards the purine ring. Finally, the ribose ring maintains a stable conformation during our simulations thanks to a hydrogen bond between C3’ -OH group and His49 side chain. This analysis probes that TGTP is firmly bound to protein, thanks to high number of intermolecular interactions.

Considering RMSD values (Figure 2B) calculated for TGTP, most residue contacts are shared among variants, except for R139C and residues Val14, Gly15, and Val16. These provide hydrophobic interactions that might be important for ligand affinity. Taking into consideration these outcomes, it seems that both the WT and the allelic variants can bind TGTP, and those differences observed in catalytic activity are not due to NUTD15 affinity towards the ligand *per se*. Due to the stability shown for the ligand during our MD simulations, it is expected that other nucleotides bind and behave in a very similar way.

After establishing the molecular recognition of TGTP by NUTD15, we turned our attention to differences in protein motions and protein intrinsically flexibility. To accomplish this, we first investigated the root mean square fluctuation during equilibrated dynamics (Figure 4 and supporting info. Figure S3).

**Figure 3.**
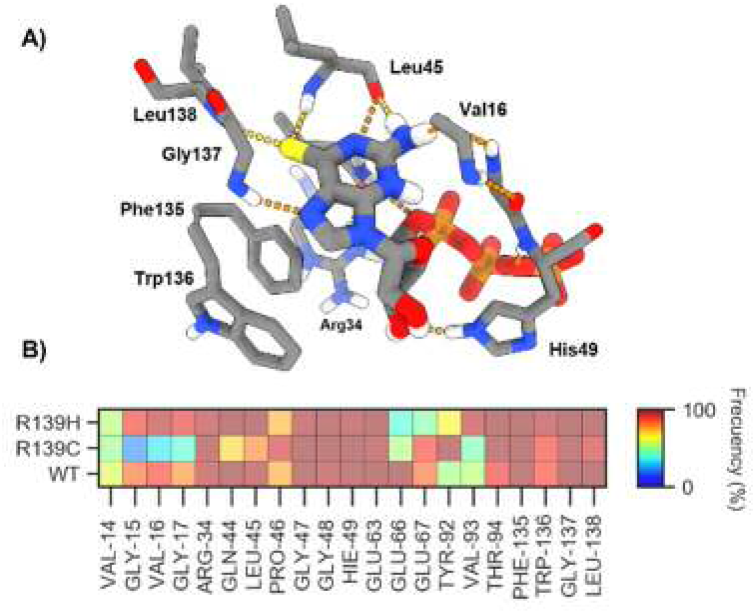
A) Representative snapshot from MD showing the main interactions between TGTP and NUTD15 residues present in the active site. B) Most frequent contacts detected during MD simulations between TGTP and active site residues.

**Figure 4.**
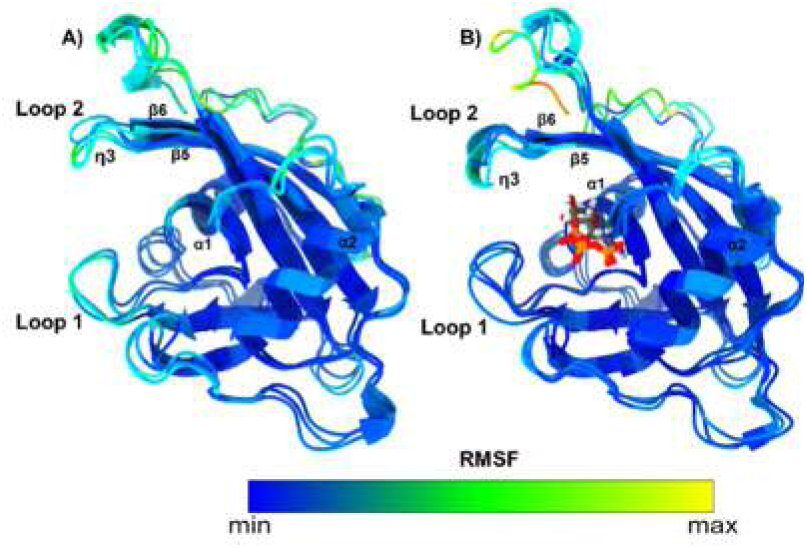
A) Representative snapshots from MD simulations of NUTD15 apo proteins superimposed and colored according to the RMSF. B) Representative snapshots from MD simulation of NUTD15 holo complexes superimposed and colored according to RMSF.

Both *C-* and *N*-termini, presented high mobility, as previously mentioned, which is expected since they lack the stabilization of the complementary NUTD15 protein of the homodimeric assembly. Additionally, this is also supported by the high temperature factors reported in the crystal structure used as a template (PDB 6T5J) (Song and others 2013). Furthermore, relatively high fluctuations were also observed in several enzyme loops. Nonetheless, interesting breakthroughs arose from RMSF calculations when comparing holo and apo states. These involve two loops that span residues Gly37-Ser42 (loop 1) and Ile85-Tyr90 (loop 2), responsible for enclosing the TGTP molecule in the catalytic throat (Figure 5A). In the initial modeled structure, loop 2 is arranged in the form of a 310-helix, known as the η3 helix. During our simulation, this portion loses partially its helicity of 310-helix, yielding an uncoiled loop (loop 2) which delimits the borders of the catalytic active site. As consequence, this motif oscillates actively and participates in TGTP occluding. Binding of the TGTP molecule leads to a partial blockage of loop 1 (residues Gly37-Ser42), which could be easily observed in the decrement of RMSF for this region (Figure 4 and supporting info Figure S2). In contrast, loop 2 remains partially immobile, with subtle fluctuations, especially in the R139C variant. This observation suggests that loop 1 plays a key role in anchoring the substrate inside the shallow active site and partially closing the catalytic throat, while loop 2 must preserve some mobility during enzyme catalysis.

**Figure 5.**
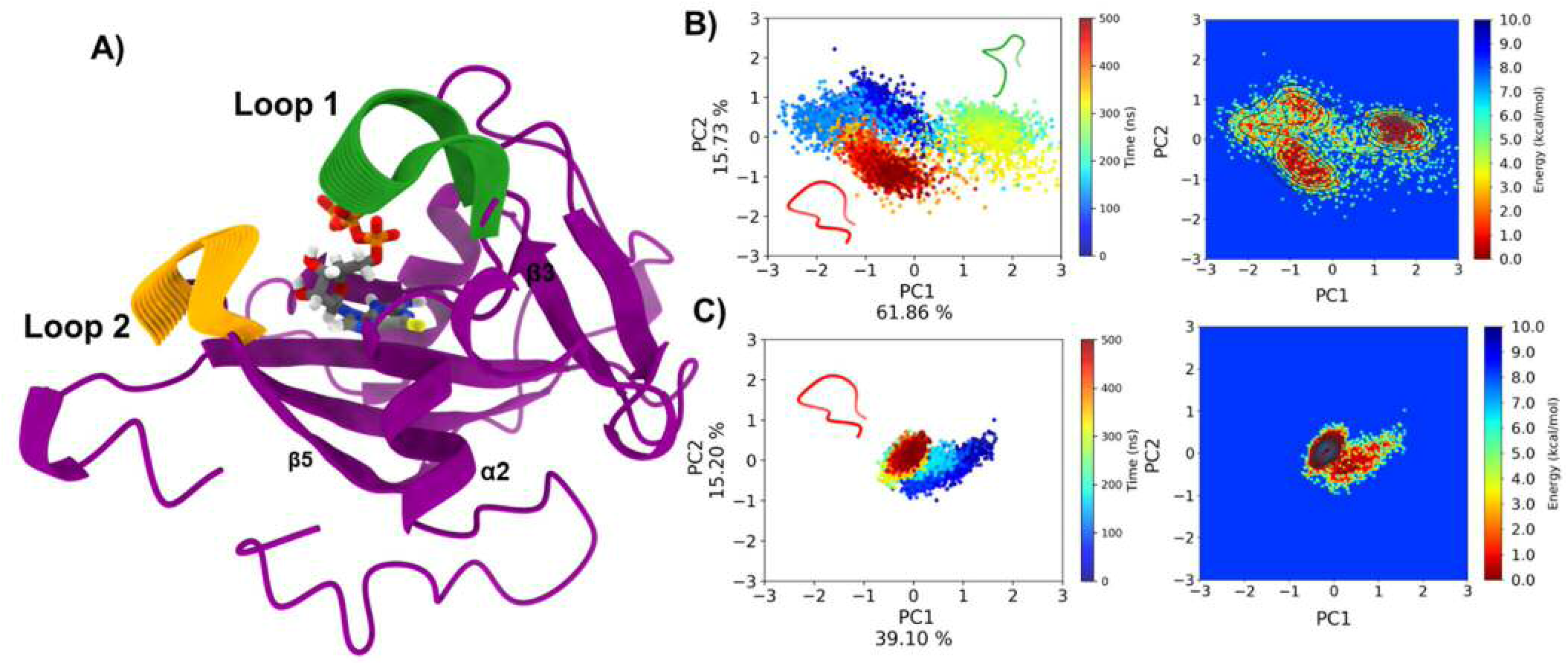
A) Representative structure from MD simulations showing loop 1 (residues Gly37-Ser42) and loop 2 (residues Ile85-Tyr90) movements identified through PCA analysis. B) On the right, projection of two PCA-calculated principal components and percentage contribution to total variance of NUTD15 WT apo with a representative structure from the MD trajectory of loop 1 in open (green) and closed (red) conformation. Left: landscapes of free energy derived from PCA projections of NUTD15 WT apo. C) On the left projection of two principal components from PCA calculations and percentage contribution to the total variance of NUDT15 WT bound to TGTP with Representative structure from MD trajectory of the loop 1 in closed conformation (red). On the right free energy landscapes determined from PCA projections of NUTD15 WT bound to TGTP.

### 2.1 Essential dynamics of NUTD15 loops

Once we discovered the different responses of these loops to ligand binding, we decided to get a deeper insight into their structural role. To fully characterize the dynamic behavior of these two partners, a Principal Component Analysis (PCA) was performed using the cartesian coordinates of backbone atoms forming in both loops (Figure 5B and 5C and supporting info. Figure S3 and S4). This type of analysis have been successfully employed to describe the essential dynamics of biomolecules and decipher the main motions of protein atoms (David and Jacobs 2014). By calculating the first five principal components, we were able to capture subtle openingclosing movements from both loops among the first and second principal components, which are responsible for the highest contributions to the system total variance (~ 35-75 %) and motions (Figure 5). This analysis corroborates the hypothesis that both loops participate actively in the structural configuration of the active site, and so, they can be part of a putative path for ligand/substrate entrance.

As above mentioned, loop 1 reduces its mobility upon TGTP binding, and this tendency is also well illustrated in the projection of the first two components, which reveals that this protein segment explores a more restricted conformational space (Figure 5 and supporting info. Figure S4 and S5). This behavior is especially notorious for the case of WT NUTD15, whose loops explore different energy minima in the configurational landscape when no nucleotide is present in the active site (Figure 5B). After ligand binding, the loop is almost confined in an energy minimum, which corresponds to a close state of the loop observed during classical MD simulations. A detailed visual inspection of loop 1 closed conformation identified during MD simulations showed that it is quite similar to that observed in crystallographic structure (PDB ID: 5LPG) (Valerie and others 2016), but even narrower, probably due to the simulation conditions mimicking a physiological-like environment.

Regarding the other two variants, it was observed that R139C mutant locates its loop 1 in a similar conformation as the WT; however, this was not observed in the case of R139H, whose loop oscillates between two different conformations according to our PCA analysis (see supporting info. Figure S4 and S5). Considering the enzymatic activity data reported in the literature (Table 1), this loop, despite being far away from phosphate reactive groups, should be quite immobile during enzyme catalysis. This scenario was not observed for the R139H model, which could be related to its lowest enzymatic activity. To further characterize the overture degree of the enzyme, we traced the distance between alpha carbons of Gly38 and Glu87, present in loop 1 and loop 2 respectively. Bidimensional distribution plots allowed us to observe that apo structures was tended to exhibit greater distribution of distances, with values near 15 Å compared to TGTP bound complexes (Figure 6A). The ligand bound complexes presented thinner peaks of distance distribution near 10 Å. These findings support our hypothesis that nucleotide binding leads to a closer conformation of the enzyme, which is mediated by these two loops.

**Figure 6.**
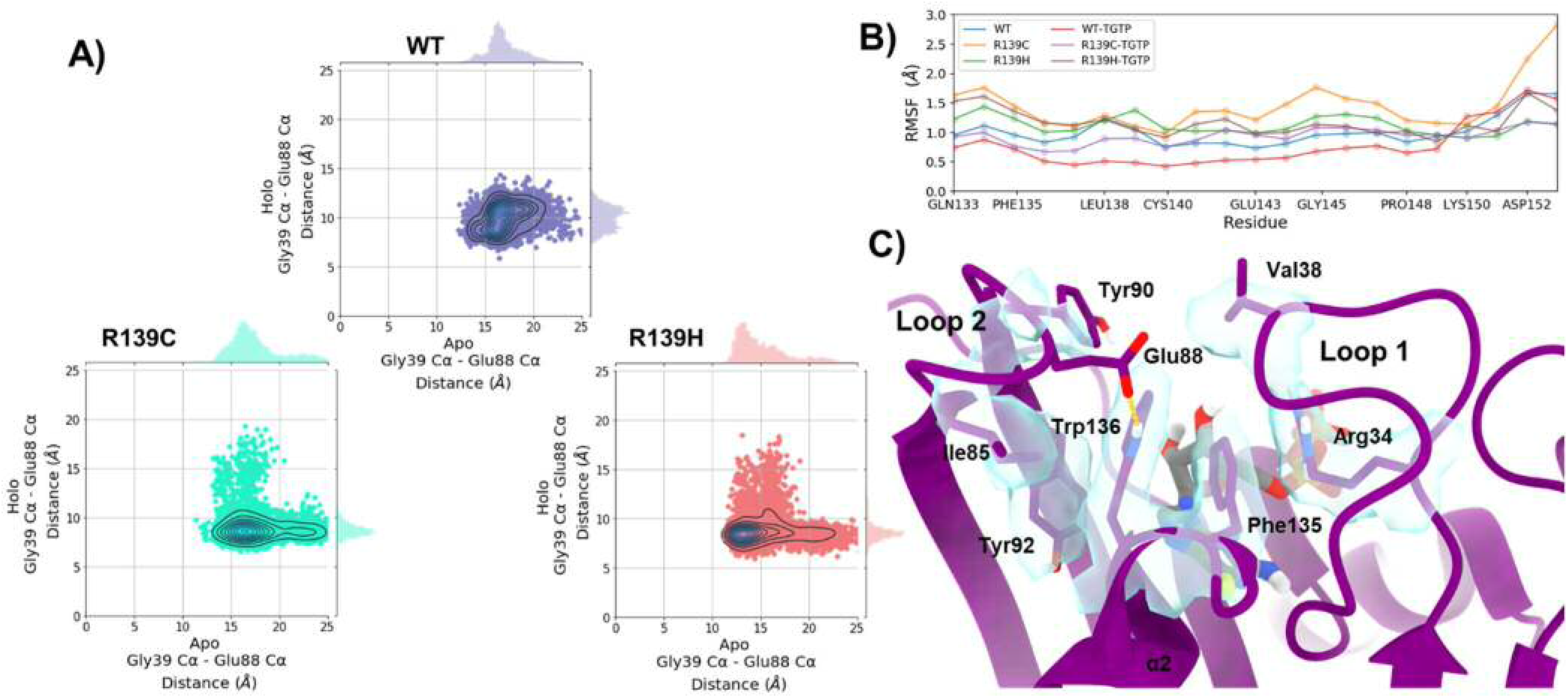
A kernel density estimate (KDE) map showing the Cα distances between Gly38 (loop 1) and Glu87 (loop 2) residues. The RMSF values for the NUTD15 α2 helix residues. B) Representative snapshots from MD simulation of NUTD15 WT in complex with TGTP showing the cluster of π-stacking interactions. Residues are embedded in a light blue isosurfaces showing their occupancy at iso level of 0.65.

Based on these discoveries, it is reasonable to hypothesize that NUTD15 must be able to close, or at least delimit, its active site cavity with these two loops in order to be a catalytically efficient enzyme. Clinically relevant variants R139C and R139H, undergoes point mutations in a residue which is located in the α2 helix. According to crystallographic reports, mutations in this solvent-exposed region produce instability and distortions in the helix (Rehling and others 2021). A more exhaustive analysis of this region let us to observe that RMSFs values are slightly increased when comparing variants with WT (Figure 6B and supporting info. Figure S3). Furthermore, this phenomenon is even more pronounced in holo structures, when the TGTP is bound in the protein active site. A mutation of Arg139 by Cys or His leads to the loss of ionic connections with Asp132, present in the α2 helix. This interaction seems to play a key role in the stability of this helix where Trp136 is located (Valerie and others 2016). However, when we monitored the hydrogen bond for the interaction between Arg139 and Asp132 side chains, we just observed a 15% and 1.84% occupancy for apo and holo systems respectively. This seems to indicate that, despite this interaction affects to the whole stability of NUTD15, its final contribution is moderate.

A closer inspection of movements and residues present in this α2 helix revealed a network of π-stacking interactions that keep NUTD15 closed across loops 1 and 2. The key point of this hub of hydrophobic interactions is the Trp136, which constitutes an anchoring point between both loops (Figure 6C). This amino acid interacts with loop 1 via CH-π interactions with Val38 -CH3 groups. On the other hand, Glu88 from loop 2 is occasionally hydrogen-bonded through its carboxylate group with the indole -NH group from Trp136. Furthermore, Phe135 also present in α2 helix, is sandwiched by Trp136 and Arg34 from loop 1 which in turn, maintains interconnected the α2 helix and loop 1 thanks to a parallel π-cation stacking interaction, as well as hydrophobic contacts with Val38. Additionally, Trp136 closes NUTD15 via CH-π interactions with Ile85 methyl groups and the aliphatic portion of the Glu86 sidechain. During MD simulations, it was also discovered that Tyr90 and Trp136 can establish parallel π-stacking interactions. In these cases, Tyr90 displaced Ile85 and Glu88, which resulted in a less compact but close protein packing. These observations are well illustrated when the percentage of contact occupancy for Trp136 and surrounding residues are studied (Table 2).

**Table 2.**
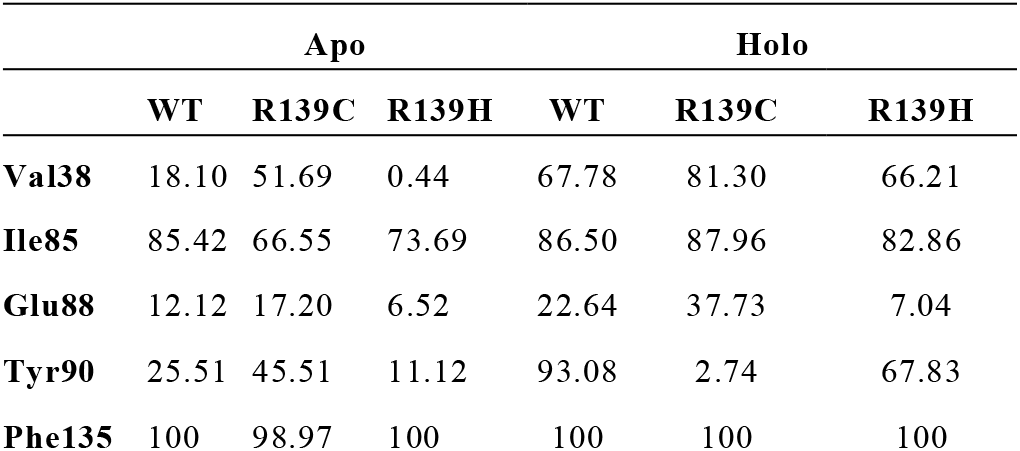
Calculated percentage of Trp136 contacts along MD simulations within a 4 Å cutoff.

|  | Apo |  |  | Holo |  |  |
| --- | --- | --- | --- | --- | --- | --- |
|  | WT | R139C | R139H | WT | R139C | R139H |
| **Val38** | 18.10 | 51.69 | 0.44 | 67.78 | 81.30 | 66.21 |
| **Ile85** | 85.42 | 66.55 | 73.69 | 86.50 | 87.96 | 82.86 |
| **Glu88** | 12.12 | 17.20 | 6.52 | 22.64 | 37.73 | 7.04 |
| **Tyr90** | 25.51 | 45.51 | 11.12 | 93.08 | 2.74 | 67.83 |
| **Phe135** | 100 | 98.97 | 100 | 100 | 100 | 100 |

The greatest change in residue contacts corresponds to Val38, which increases its contacts with Trp136 upon ligand binding, being especially notorious for WT and R139H. This is the main connector between loop 1 and α2 helix thanks to the hydrophobic and CH-π interactions with the Trp136 indole ring. Those contacts observed on Glu88 are mainly due to a hydrogen bond with -NH from Trp136, as above mentioned, when the nucleotide is present in the active site. This interaction represents a direct path to enclose loop 2 towards loop 1. Therefore, occasional hydrophobic interactions could be found between the aliphatic portion of the Glu88 side chain Ile85 and Trp136. Finally, Phe135, contacts are the most prevalent due to its close proximity to Trp136 in order to establish parallel the π-stacking. The stabile contact between Phe135 and Trp136 correspond to high prevalent parallel π-stacking, which transmits movements in the α2 helix.

Based on these findings, it appears that situations that cause Trp136 fluctuation may disrupt this cluster of residues and their interactions, altering the functional structure of the enzyme. Indeed, RMSF values for Trp136 in R139C (0.88 Å) and R139H (1.21 Å) mutants with TGTP bound, are higher when comparting with WT (0.5 Å) NUTD15. Thus, instability in this molecular anchor leads to fluctuations in the whole architecture of the active site region, especially in loops 1 and 2. This is in very good agreement with relative activities reported in the literature (Table 1) for *in vitro* assays, supporting our hypothesis (Rehling and others 2021). Moreover, the importance of Trp136 as a structural node in NUTD15 is also supported by the small molecule inhibitor TH7755 which binding mode is stabilized by hydrophobic interactions with Val38 (Rehling and others 2021). Consequently, the design of chemical probes targeting this residue could yield NUTD15 inhibitors a mechanism independent of substrate active site competition.

## 4 Conclusions

The investigation of the dynamical behavior of NUTD15 and some of its most important clinical variants with atomic detail, allowed us to further understand some structural features and protein functions not previously observed or described.

Our study highlights the strong contribution of the TGTP nucleotide to the conformational stability of the whole protein once bound, which is consistent with evidence for WT and R139C variant. In addition, the conformational landscape of two loops has been investigated in detail. In the absence of any bound nucleotide loop 1 (Gly37-Ser42) and loop 2 (Ile85-Tyr90) display moderate fluctuations. Upon ligation of TGTP nucleotide, loop 2 must be partially blocked, while loop 1 must retains some mobility to be NUTD15 catalytically efficient. As such, it seems that these loops can serve as gatekeepers harboring for the nucleotide. Our PCA analysis revealed that this loop spans a larger conformational landscape during MD simulations for the R139H point variation. Furthermore, thanks to a network of hydrophobic contacts mediated by Trp136, these protein segments are important for maintaining NUTD15 in a compact shape, possibly critical for its catalytic mechanism of action. This residue present substantial fluctuations in R139C and R139H variants, destabilizing the network of hydrophobic and π-interactions, which in turn, might account for the diminished NUTD15 enzymatic activity. As such, it appears that mutation of residues present in α2 helix can affect to the architecture of this protein by perturbing mobility of these loops.

Our investigation not only let us gain insight into the experimental data of this hydrolase but also illustrated how point mutations affect the conformational diversity of this protein, affecting its enzyme activity. This knowledge may be of interest for the understanding of new point mutation variants, as well as for the design of new inhibitors targeting a possibly non-competitive mechanism of action that leads to the blockage of NUTD15 loops.

## Acknowledgements

Javier Garcia-Marin thanks to the Spanish Ministry of Universities and University of Alcala for the postdoctoral fellowship ‘Margarita Salas’.

## Authorship contribution statement

Elena Gomez: Calculations, analysis and manuscript review. Javier Garcia-Marin: Conceptualization, calculation, analysis, writing - original draft, review and editing.

## Abbreviations

RMSD: Root Mean Square Deviation
RMSF: Root Mean Square Fluctuation
MD: Molecular Dynamics
PCA: Principal Component Analysis.

